# Dorsomedial prefrontal cortex activity during learning discriminates response to Cognitive Behavioural Therapy in depression

**DOI:** 10.1101/224410

**Authors:** Filippo Queirazza, Elsa Fouragnan, J. Douglas Steele, Jonathan Cavanagh, Marios G. Philiastides

## Abstract

**BACKGROUND:** Cognitive behavioural therapy (CBT) is an effective evidence based treatment for depression. At present there is no reliable predictor of CBT in depression. Although the key to successful CBT in depression lies in altering maladaptive information processing, no previous imaging study has probed predictors of CBT response using pre-treatment neural encoding of information processing. METHODS: Using functional magnetic resonance imaging we scanned 37 unmedicated depressed subjects before and after completing computerised CBT (cCBT). We model the trial-by-trial appraisal of feedback information during a probabilistic learning task by means of a dynamic learning rate. To discriminate response to cCBT we capitalise on the pre-treatment blood oxygen level dependent (BOLD) activity encoding the dynamic learning rate as a function of feedback congruence and valence. Additionally, we probe between-group differences in the learning style encoded in the model’s parameters.

**RESULTS:** We show BOLD activity in the dorsomedial prefrontal cortex (dmPFC) to be encoding the dynamic learning rate. Crucially, responders exhibit greater BOLD activity in the dmPFC during incongruent negative trials but lower BOLD activity during congruent negative trials than non-responders. Additionally, on between-group comparisons of model’s parameter estimates we show responders take relatively greater account of previous feedback history and make comparatively smaller adjustments to the learning rate as a result of outcome surprisingness.

**CONCLUSIONS:** Our findings provide novel and important insights into the cognitive mechanisms underpinning response to cCBT and lend support to the feasibility and validity of neurocomputational approaches to treatment prediction research in psychiatry.

## INTRODUCTION

The cognitive model of depression posits that biased acquisition and processing of feedback information gives rise to and perpetuates depressive symptoms (1, 2). Based on this theoretical formulation, Aaron T. Beck developed Cognitive Behavioural therapy (CBT) as a treatment for depression (3). CBT is an effective evidence-based intervention for depressive disorder (4, 5). In the UK computerised CBT (cCBT) (that is, self-help internet-delivered CBT) is recommended as a treatment option for mild to moderate depression in the National Institute for Health and Care Excellence guidelines.

Despite recent multidisciplinary efforts (6-12), there is no reliable predictor of clinical response to CBT in depression (13, 14). Neural predictors using functional magnetic resonance imaging (fMRI) provide the additional advantage of illuminating the functional neuroanatomy that underpins response to CBT in depression (15).

Remarkably, although the clinical practice of CBT in depression primarily involves evaluating and ultimately correcting negatively biased inferences drawn from probabilistic information, only emotion eliciting (6, 16, 17) or task-free resting-state paradigms (8, 18, 19) have so far been employed to probe pre-treatment fMRI predictors of CBT response.

In contrast, the computational framework of value-based learning (also known as reinforcement learning) paradigms affords a rigorous and quantitative account of the cognitive mechanisms implicated in drawing inferences from probabilistic feedback. Crucially, within this framework, a weighting factor known as the learning rate controls how the value of unexpected feedback information (also know as the prediction error) is appraised. (20). In experimental settings mimicking real-life, volatile environments, adaptive value-based learning is implemented via a dynamic (that is, time varying) rather than constant learning rate (21-25). Abnormal tuning of the dynamic learning rate distorts processing of probabilistic feedback and impairs learning (26). Prior behavioural evidence suggests that tuning of the dynamic learning rate is disrupted in depression as a function of feedback congruence and valence (27, 28).

In this work, we capitalise on the neural encoding of the dynamic learning rate during probabilistic learning to discriminate response to cCBT. Since successful CBT involves cognitive restructuring of negatively biased thinking we predicted processing of probabilistic negative feedback would uncover between-group differential pre-treatment BOLD activity. Additionally, we found that a learning style that facilitates reframing of negative cognitive biases was associated with response to cCBT.

## METHODS AND MATERIALS

### Sample

All 37 participants (18 women) were recruited via self-referral through local newspaper advertisement. Eligibility criteria were a primary diagnosis of depressive disorder as operationalised by ICD-10 diagnostic criteria and a score ≥ 14 on the Beck’s Depression Inventory-II (BDI-II). To avoid any potential confound associated with psychotropic medications, we only recruited unmedicated depressed subjects. Exclusion criteria included current involvement with other CBT-based interventions or psychological therapies, a comorbid diagnosis of other major mental disorder, CBT treatment within past 3 years, a diagnosis of psychoactive substance dependence, previous history of brain injury.

Eleven subjects (~29.7%) did not attend the post-treatment assessment. Of these subjects, 6 deteriorated and required treatment with antidepressant medications and 5 did not complete cCBT due to lack of efficacy. High dropout rates are common in studies examining Internet delivered psychotherapies (12). Twenty-six subjects (~70.3%) attended the post-treatment appointment and of these only one was unable to undergo scanning.

In total 19 subjects were classified as responders and 18 subjects were classified as non-responders. The overall cCBT response rate was 51.3%. The average post-treatment improvement in BDI-II score was around 62% (± 40%).

Between-group comparisons revealed no significant differences in age (t_35_=0.08; p=0.93) and sex (χ^2^_1_=0.02; p=0.86). Non-responders had a significantly higher pre treatment BDI-II score than responders (t_35_=2.86; p=0.006). On further sensitivity analyses we found this significant difference to be mainly determined by those non-responders who did not complete cCBT (t_27_._3_=4.74; p<0.001) rather than by those who did (t_24_=0.62; p=0.53). In spite of these between-group differences in pre-treatment BDI-II scores, we wanted to probe differences in the neurocomputational mechanisms underlying probabilistic learning in order to expose the internal cognitive mechanisms implicated in response to cCBT.

All participants provided written, informed consent. The study protocol was approved by the West of Scotland Ethics Committee (10/S0703/71).

### Experimental design

This study adopted a naturalistic longitudinal design. Participants attended two appointments (pre- and two months post-treatment with cCBT) (see Supplemental Materials for further details on cCBT and study design). Each appointment included a clinical evaluation by a qualified psychiatrist followed by an fMRI scan. A clinical diagnosis of depression was corroborated using the Clinical Interview Schedule – Revised (CIS-R) (29). To measure depression severity we used the BDI-II, which is a clinically-validated tool to assess intensity of depression (30).

We regarded non-completion of cCBT as an index of treatment failure and classified all non-completers as non-responders. For the remaining subjects response to cCBT was defined as a 50% or greater reduction in the pre-treatment BDI-II score. To account for response bias we performed sensitivity analyses excluding non-completers from the non-responders group.

### fMRI experimental paradigm

To probe the neural correlates of probabilistic reinforcement learning we employed a probabilistic reversal-learning task during fMRI (Figure 1). This task involved learning which of two stimuli yielded the highest payoff rate and included 180 trials lasting approximately 20 minutes. Participants were required to choose between two abstract visual stimuli both yielding either positive (+10) or negative (-10) outcome devoid of monetary value. Stimulus-outcome contingencies were probabilistic and asymmetrically skewed (70–30%) (Figure 1).

**Figure 1.**
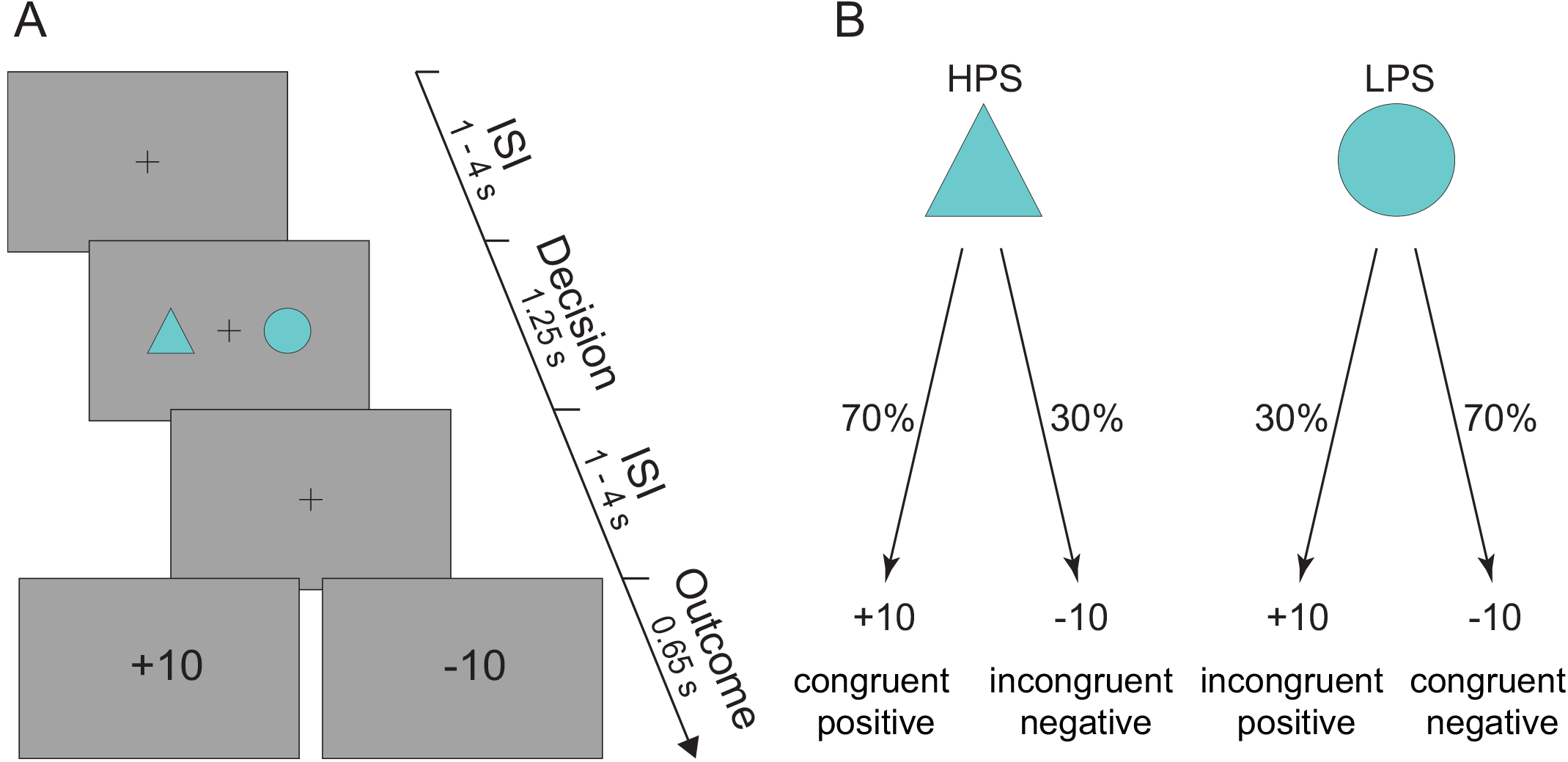
Probabilistic reversal-learning task. (A) Each trial commenced with a jittered interstimulus interval (1–4s) displaying a fixation cross. Subsequent to this two abstract visual stimuli appeared randomly on either side of the screen for 1.25s. For each participant the two stimuli were randomly chosen from a pool of 18 different geometrical shapes. Participants were given 1s to choose a stimulus via a button press. Following a second jittered interstimulus interval (1–4s) participants were presented with the outcome of their decision for 0.65s. Outcome was either positive (+10) or negative (-10). To maximize design efficiency the duration of jittered interstimulus intervals was optimized implementing a genetic algorithm described in (49). ISI: Interstimulus Interval. (B) Classification of probabilistic feedback as a function of congruence and valence. Stimulus-outcome contingencies were asymmetrically skewed (70–30%) so that the expected value of the two stimuli was of the same magnitude but of opposite sign. This meant that whilst one stimulus (here referred to as the high probability stimulus) was associated with a greater likelihood of positive outcome, the other stimulus (here referred to as the low probability stimulus) was associated with a greater likelihood of negative outcome. Reversals were self-paced and occurred when participants chose the high probability stimulus five times over the last six trials. To prevent participants from figuring out the underlying reversal rule we ran a randomly generated number of buffer trials from a zero-truncated Poisson distribution before reversing stimulus-outcome contingencies. The stimulus-outcome association strength was chosen to enable detection of reversals. Participants were only advised of the probabilistic nature of the task and that stimulus-outcome contingencies might reverse based upon their performance. HPS: High Probability Stimulus. LPS: Low Probability Stimulus.

Furthermore, to create a volatile environment stimulus-outcome contingencies were reversed in the course of the experiment. Notably, reversals were triggered when participants chose the high probability stimulus (that is, the stimulus with a greater chance of yielding a positive outcome) five times over the last six trials. As a result participants experienced a different number of reversals.

Participants were advised of the probabilistic nature of the task and that stimulus-outcome contingencies might reverse based upon their performance. Moreover, to ensure participants understood the nature of the task they underwent a 5 minutes practice session prior to the fMRI scan. The task was programmed using Presentation® (Neurobehavioural Systems) stimulus delivery software.

The probabilistic reversal-learning task is ideally suited to exposing information processing biases during probabilistic learning. Ultimately optimal performance rests upon attaching a greater weight (via a greater learning rate) to more informative *congruent* feedback (that is, feedback that is consistent with the most likely outcome associated with the chosen stimulus) than to less informative *incongruent* feedback (that is, feedback that is not consistent with the most likely outcome associated with the chosen stimulus). To do so subjects need to be able to infer whether fluctuations in observed stimulus-outcome associations reflect either noise (given underlying stochastic contingencies) or sudden environmental changes (that is, reversals).

In depression there is substantial behavioural evidence that both congruence (27, 28) and the affective quality (i.e. valence) (2) of probabilistic feedback significantly contribute to biasing information processing. Accordingly, in our analyses we leverage the combined effect of congruence and valence on the dynamic learning rate to probe behavioural and neural differences between responders and non-responders.

### Behavioural statistical analysis

To verify that individual performance was better than chance we used a binomial test to compare the number of correct choices with chance level.

To examine for between-group differences in task performance we regressed subject-wise percentage of high probability stimulus choices on clinical outcome after adjusting for pre-treatment BDI-II score. Additionally we tested for any between-group difference in the effect of the high probability stimulus on choice behaviour (the higher the effect of the high probability stimulus the more optimal the choice behaviour and thus learning) using a generalised mixed-effects linear model (see Supplemental Materials for further details on this analysis).

Finally we tested for any between-group difference in the model-derived dynamic learning rate estimates as a function of congruence and valence (see Supplemental Materials for further details on this analysis).

### Computational modelling of behavioural data

We fitted 5 different models (see Supplementary Materials for further details) that use trial-wise scaling of the prediction error but make different assumptions on the computational mechanisms supporting this dynamic tuning of learning rate.

In the winning model the learning rate scales with the slope of the smoothed unsigned prediction error (i.e. absolute value of prediction error) (24). This model reprises Pearce-Hall’s theory that surprise (formalised as the unsigned prediction error) drives the acquisition of stochastic stimulus-outcome contingencies (31) but with some important refinements. Indeed, compared to the Pearce-Hall’s model, the smoothing of the unsigned prediction error (the degree of which is regulated by a free parameter *ρ*) should render the inference process about whether a change has occurred in the environment more robust to the inherent task stochasticity. Moreover, an additional free parameter *γ* controls the extent to which the dynamic updating of the learning rate is influenced by the slope. For example, whilst lower values of *γ* yield substantial trial-by-trial changes of the dynamic learning rate even in the presence of small slope estimates (that is, low surprise), higher values of *γ* result in a more stable learning rate even in the presence of significant slope estimates (that is, high surprise). Hence, this model also allows for the possibility that subjects might be employing a relatively fixed learning rate. The decision function for all models was a standard sigmoid function parameterised by the inverse of the temperature parameter, *β*.

### Model fitting and model comparison

To optimise each model’s free parameters we implemented the hierarchical type II maximum likelihood fitting procedure described in (32) (see Supplemental Materials for further details). The fits of all models were compared using the Integrated Bayesian Information Criterion (BIC_int_) (32) (see Supplemental Materials for further details).

To ascertain any between-group differences in the learning style encoded in the model’s fixed parameters we regressed subject-wise parameter estimates against clinical outcome (response vs. non-response) after adjusting for pre-treatment BDI-II score. To account for the response bias we performed additional sensitivity analyses.

Finally we ran sanity checks on the winning model’s goodness of fit. To verify accuracy of model’s fit we first binned predicted choice propensities according to their quintiles and subsequently measured the strength of their linear association with corresponding observed choice probabilities using Pearson’s correlation coefficient. Additionally, we employed a binomial test to test whether the number of choices correctly predicted by the model exceeded that expected by chance (33).

### fMRI data acquisition

We used a 3T GE system with an 8-channel parallel imaging head coil. We acquired a high-resolution *T*_1_-weighted structural image (0.5 x 0.5 x 1 mm voxels, 320 x 320 matrix, 160 axial slices, TI = 500 ms, TR = 7700 ms, TE = 1.5 ms, flip angle = 12^o^) using an optimized Inversion Recovery Fast SPoiled GRadient echo sequence (IR-FSPGR) and a functional echo planar imaging (EPI) scan (3 mm isotropic voxels, 64 x 64 matrix, 608 axial slices, TR = 2000 ms, TE = 30 ms, flip angle = 80^o^). Slice orientation was tilted −20^o^ from the AC-PC plane to alleviate signal drop out in the orbitofrontal cortex (34). The first four volumes of the functional scan were discarded in order to allow for the magnetic field to reach the steady state.

### fMRI data preprocessing and statistical analysis

Pre-treatment and post-treatment fMRI data preprocessing and statistical analyses were performed using FSL software (35). Preprocessing pipeline involved intra-modal motion correction using MCFLIRT (36), slice timing correction, spatial smoothing with an isotropic 5 mm FWHM Gaussian kernel, high-pass temporal filtering with 110 sec. cut-off frequency and grand-mean intensity normalisation of each entire 4D dataset. Functional scans were subsequently co-registered with skull-stripped structural images using boundary-based registration (FLIRT) (37, 38) and spatially normalised into MNI152 space using FNIRT non-linear registration.

Whole brain statistical analyses of pre-treatment fMRI data were performed using a multilevel mixed-effects approach as implemented in FLAME1 (FSL) (39). At the first-level the subject-specific general linear model included the model-derived dynamic learning rate as the regressor of interest plus a number of additional nuisance covariates (see Supplementary Materials for further details). Most importantly, we included a nuisance regressor accounting for surprise (here formalised as unsigned prediction error) to retrieve the unique effect of the dynamic learning rate on BOLD activity.

To improve efficiency of our fMRI statistical analysis all model-derived regressors were obtained by generating subject-wise model fits using the population-level parameter means (40). All regressors were convolved with a hemodynamic response function (double gamma function).

We estimated the subject-wise linear contrasts of parameter estimates and subsequently entered these contrast images into a second-level mixed-effects analysis where we tested both mean group effect and between-group (responders vs. non-responders) differences. We thresholded the resulting Z statistic images using cluster-defining threshold of Z>3.1 and a FWE-corrected significance threshold of p=0.05.

Furthermore, we examined for any between-group difference in the pre-treatment BOLD activity encoding the dynamic learning rate. To this end we first retrieved BOLD percent signal change time-locked to the onset of the outcome phase from a cluster broadly corresponding to the dmPFC (max Z=7.25; −4 −2 56). We chose this cluster since it is consistent with previous reports on the fMRI correlates of the dynamic learning rate (24, 41). Subsequently we performed between-group comparisons as a function of feedback congruence and valence using mixed-effects linear models (see Supplemental Materials for further details on this analysis).

We also assessed for a robust linear association between subject-wise (that is, aggregated up to the subject-level) pre-treatment event-locked BOLD percent signal change and post-treatment symptomatic improvement.

Having ascertained between-group differences in pre-treatment BOLD activity we sought to determine whether such BOLD activity was either a moderator or mediator of cCBT response. (see Supplemental Materials for further details on this analysis).

## RESULTS

### Clinical outcome

Between-group comparisons revealed no significant differences in age (t_35_=0.08; p=0.93) and sex (χ^2^_1_=0.02; p=0.86) between responders and non-responders. Non-responders had a significantly higher pre-treatment BDI-II score than responders (t_35_=2.86; p=0.006). On further sensitivity analyses we found this significant difference to be mainly determined by those non-responders who did not complete cCBT (t_27_._3_=4.74; p<0.001) rather than by those who did (t_24_=0.62; p=0.53). In spite of these between-group differences in pre-treatment BDI-II scores, we wanted to probe differences in the neurocomputational mechanisms underlying probabilistic learning in order to expose the internal cognitive mechanisms implicated in response to cCBT.

### Task performance and behavioural estimates of the dynamic learning rate do not discriminate response to cCBT

All participants did not choose randomly (binomial test, mu=0.5, p<0.05) during the task. We did not find any significant between-group difference in the number of buffer trials (t_35_=-0.41; p=0.68) during the pre-treatment experiment. Task performance was similar across responders and non-responders (see Supplemental Results for further details).

Pre-treatment behavioural estimates of the dynamic learning rate as a function of congruence and valence did not discriminate between responders and non-responders (congruent positive: t_38.77_=-0.24, p=0.80; incongruent positive: t_37.2_=0.45, p=0.65; congruent negative: t_38_._07_=0.30, p=0.76; incongruent negative: t_159_._97_=-0.41, p=0.67) (Figure 2).

**Figure 2.**
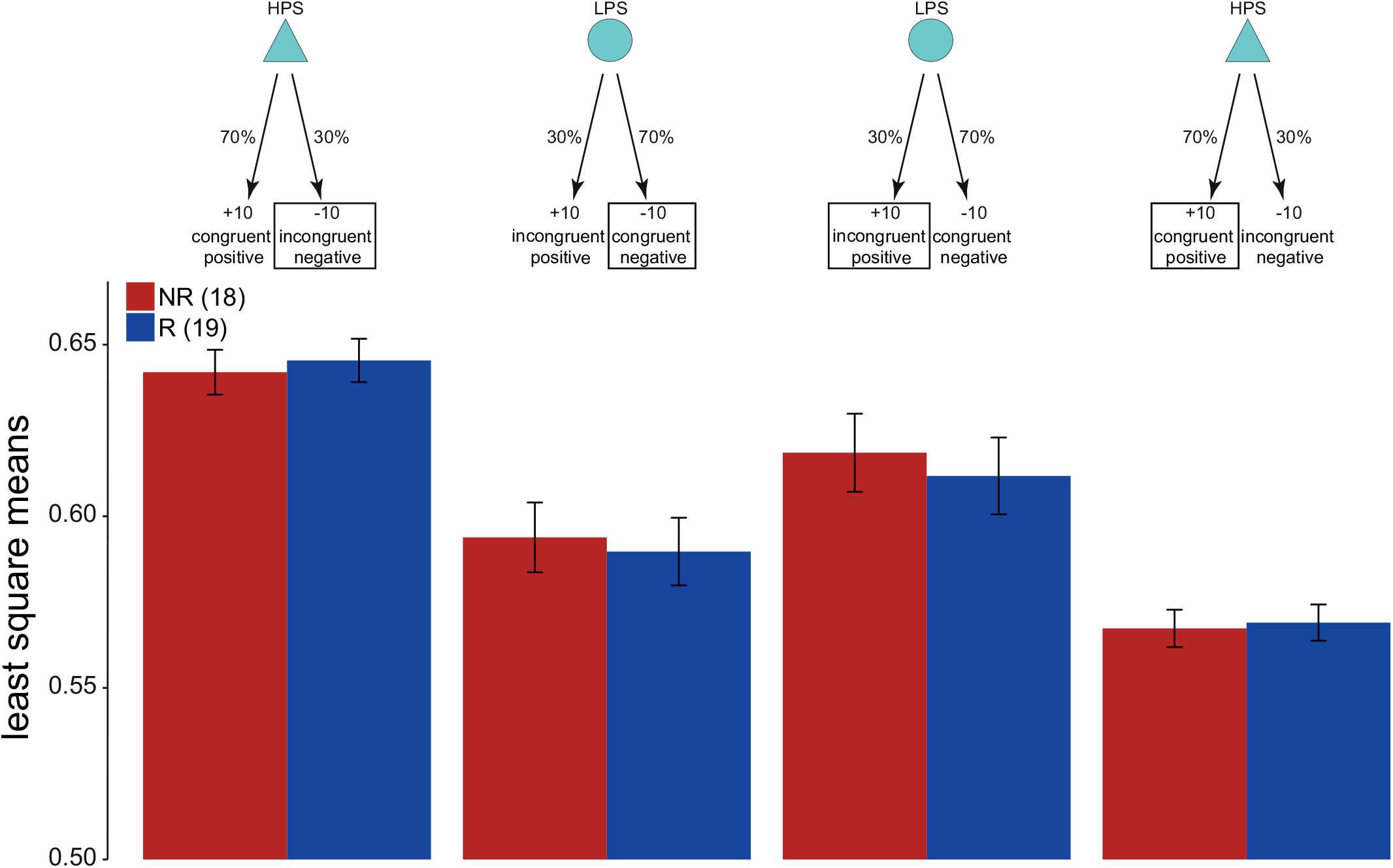
Analysis of behavioural estimates of dynamic learning rate. Least-square mean estimates ± SEM (error bars) of dynamic learning rate in the responders (blue; n=19) and non-responders (red; n=18) groups are plotted as a function of feedback congruence and valence. None of the post-hoc between-group comparisons were statistically significant.

### Model comparison

On formal Bayesian model comparison we found that the best fitting model was that described in (24) with a BIC_int_ of ~287 (32). (Krugel et al. model with additional parameter *α*^1^: BIC_int_=540.82; Hierarchical Gaussian Filter: BIC_int_=828.68; Pearce-Hall: BIC_int_=332.72; Kalman filter K1 variant: BIC_int_=708.70) (Figure 3A).

**Figure 3.**
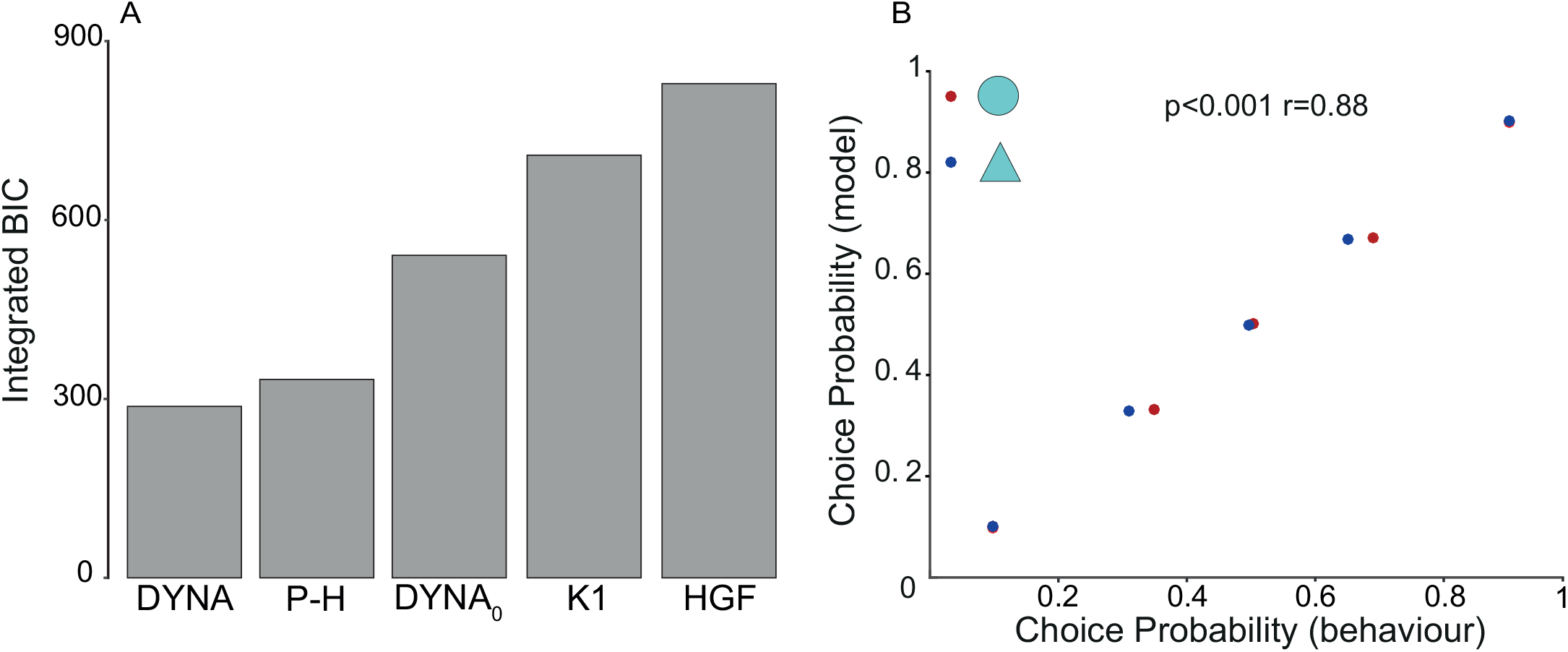
Computational model comparison and behavioural fit. (A) BICint scores for all models. Lower scores indicate better fit. DYNA is the winning model. DYNA: Krugel et al.’s model. DYNA_0_: Krugel et al.’s model with additional parameter *α*^1^. HGF: Hierarchical Gaussian Filter. P-H: Pearce-Hall. K1: Kalman filter K1 variant. (B) Scatterplot shows linear relationship between empirical and predicted choice probabilities. r: Pearson’s correlation coefficient.

We verified the model’s goodness of fit using a binomial test and found that under the null hypothesis that on each trial the model was choosing at chance level, the probability of model’s correctly predicted *n* choices was <0.05 across all subjects. Additionally we found that observed and model’s predicted choice probabilities were significantly correlated (r = 0.88; p<0.001), further endorsing the quality of the model fits (Figure 3B).

### Parameters reflecting specific information processing style are associated with differential response to cCBT

After adjusting for pre-treatment BDI-II score, we found a statistically significant between-group difference in the estimates of the model’s parameter *ρ* (logit(*ρ*): t_34_=-2.15; p=0.038) and *γ* (log(*γ*): t_34_=2.11; p=0.041) (Figure 4) but no significant difference for *β* (log(*β*): t_34_=0.61; p=0.54). Additional sensitivity analyses revealed these significant differences to be predominantly driven by non-responders who did not complete cCBT (logit(*ρ*): t_27_=-3.12; p=0.004; log(*γ*): t_27_=2.99; p=0.005) rather than by those who did (logit(*ρ*): t_23_=-0.72; p=0.47; log(*γ*): t_23_=0.65; p=0.51).

**Figure 4.**
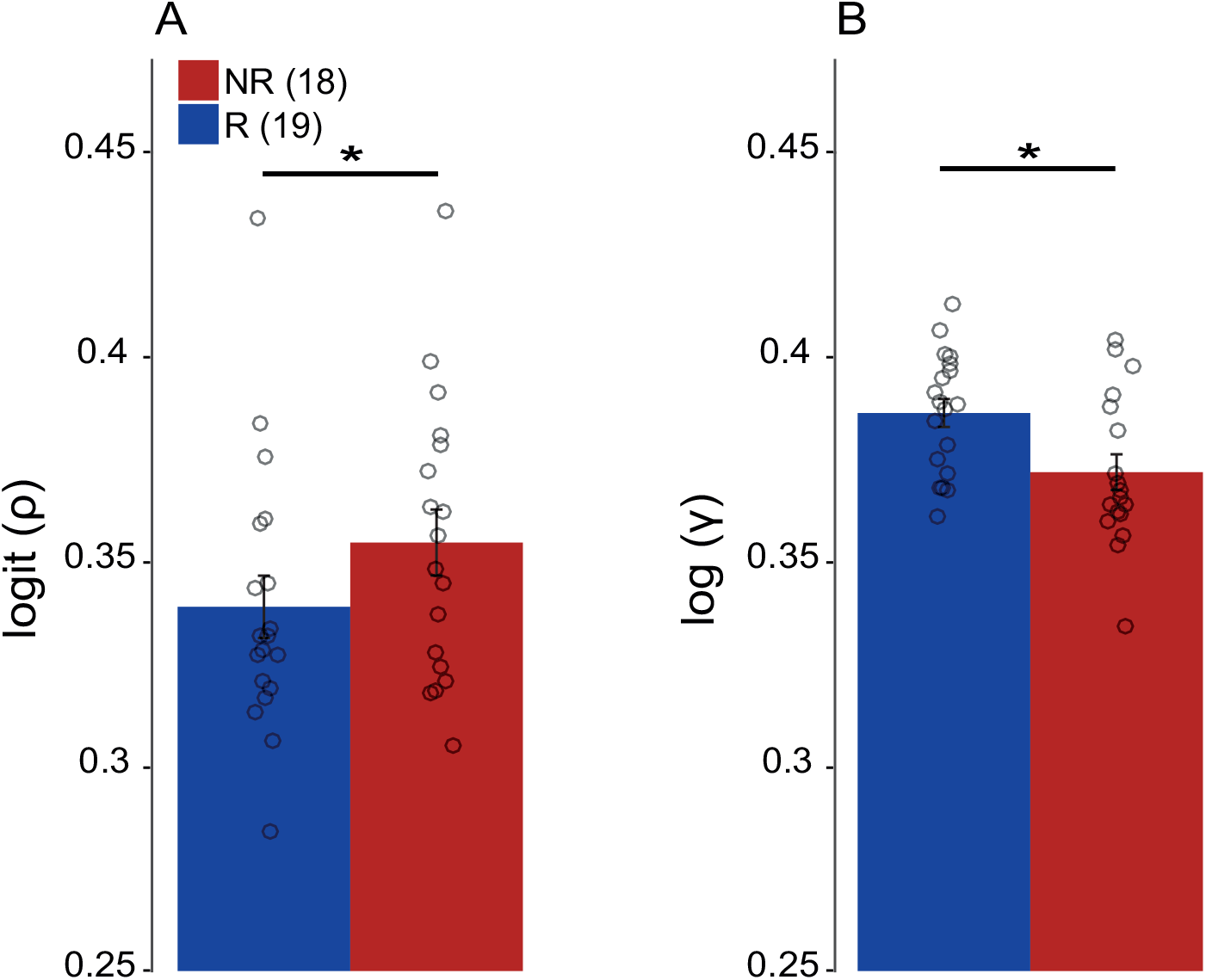
Computational parameters. Mean estimates ± SEM (error bars) of *ρ* (left) and *γ* (right) in the responders (blue) and non-responders (red) groups. Parameter estimates are shown in their native space (logit for *ρ* and log for *γ*). After adjusting for pre-treatment BDI-II score, on average responders exhibited a greater tendency to smooth over previous unsigned prediction errors (lower *ρ* mean estimate) and to make smaller adjustments to the dynamic learning rate (greater *γ* mean estimate) compared to non-responders. Black circles represent individual subjects. * p < 0.05.

In sum, responders were on average more prone (that is, lower mean estimate of *ρ*) to smoothing over previous unsigned prediction errors than non-responders. This implies that responders took greater account of previous feedback history than non-responders. Additionally, on average responders had a tendency to make relatively smaller trial-wise adjustments of the learning rate (that is, greater mean estimate of *γ*) as a result of outcome surprisingness than non-responders.

### Neural signature of dynamic learning rate

Whole-brain model-based fMRI analysis of pre-treatment fMRI data revealed significant bilateral activations correlating with the dynamic learning rate in a set of brain regions broadly located in the dorsomedial prefrontal cortex (dmPFC), dorsolateral prefrontal cortex (including precentral gyrus, postcentral gyrus) and occipital cortex (fusiform gyrus, lingual gyrus) (Supplemental Table S1). Significant activations survived family-wise error (FWE) correction for multiple comparisons (p < 0.05) at the cluster level with a cluster-defining threshold of p < 0.001.

A number of previous studies have suggested that the dmPFC in particular might be involved in the dynamic online adjusting of the learning rate (22, 41, 42), although they employed different samples, experimental paradigms and modelling assumptions. Moreover, previous work reported abnormal activation of this region in response to probabilistic feedback in a sample of unmedicated depressed subjects compared to healthy controls (28). We therefore used BOLD activity in the bilateral dmPFC shown in Figure 5 to discriminate treatment response (see Materials and Methods for further details).

**Figure 5.**
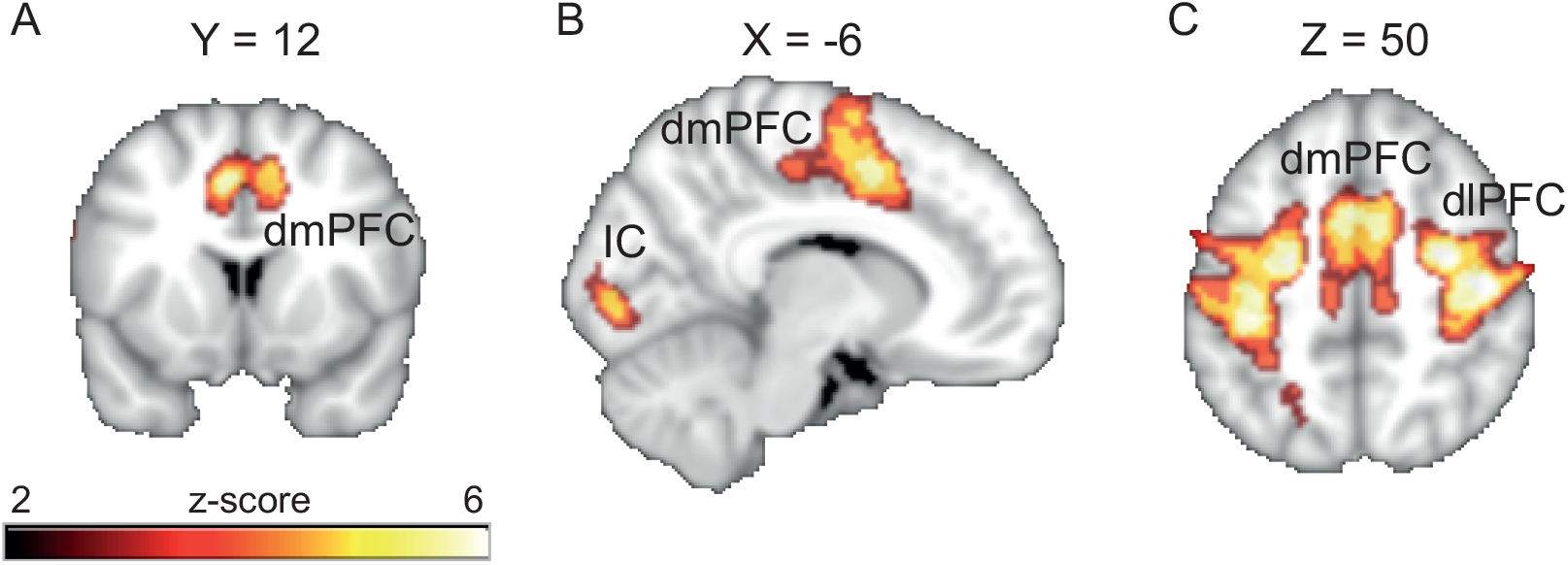
Pre-treatment whole-brain results for dynamic learning rate contrast. Activations shown here survived a cluster-defining threshold of Z > 3.1 and FWE-corrected significance threshold of p = 0.05. (A) dmPFC: dorsomedial prefrontal cortex. (B) IC: intracalcarine cortex (C) dlPFC: dorsolateral prefrontal cortex. Coordinates are given in the MNI space. We used the cluster in the dmPFC to extract trial-wise feedback-locked BOLD percent signal change for all participants and used them to discriminate treatment response.

### Pre-treatment BOLD activity encoding dynamic learning rate discriminates between responders and non-responders

A two-sample unpaired t-test on the dynamic learning rate contrast images did not identify any significant between group differences. However, on between-group comparisons of the outcome-locked BOLD activity in the dmPFC we found that responders exhibited significantly greater BOLD percent signal change during incongruent negative trials (F_1,34.07_=8.87, p=0.005) and significantly lower BOLD percent signal change during congruent negative trials (F_1,33.01_=5.15, p=0.029) compared to non-responders (Figure 6). No significant between-group differences were found following positive feedback (congruent: F_1,34.72_=1.45, p=0.23; incongruent: F_1,33.79_=0.33, p=0.56) (Figure 6).

**Figure 6.**
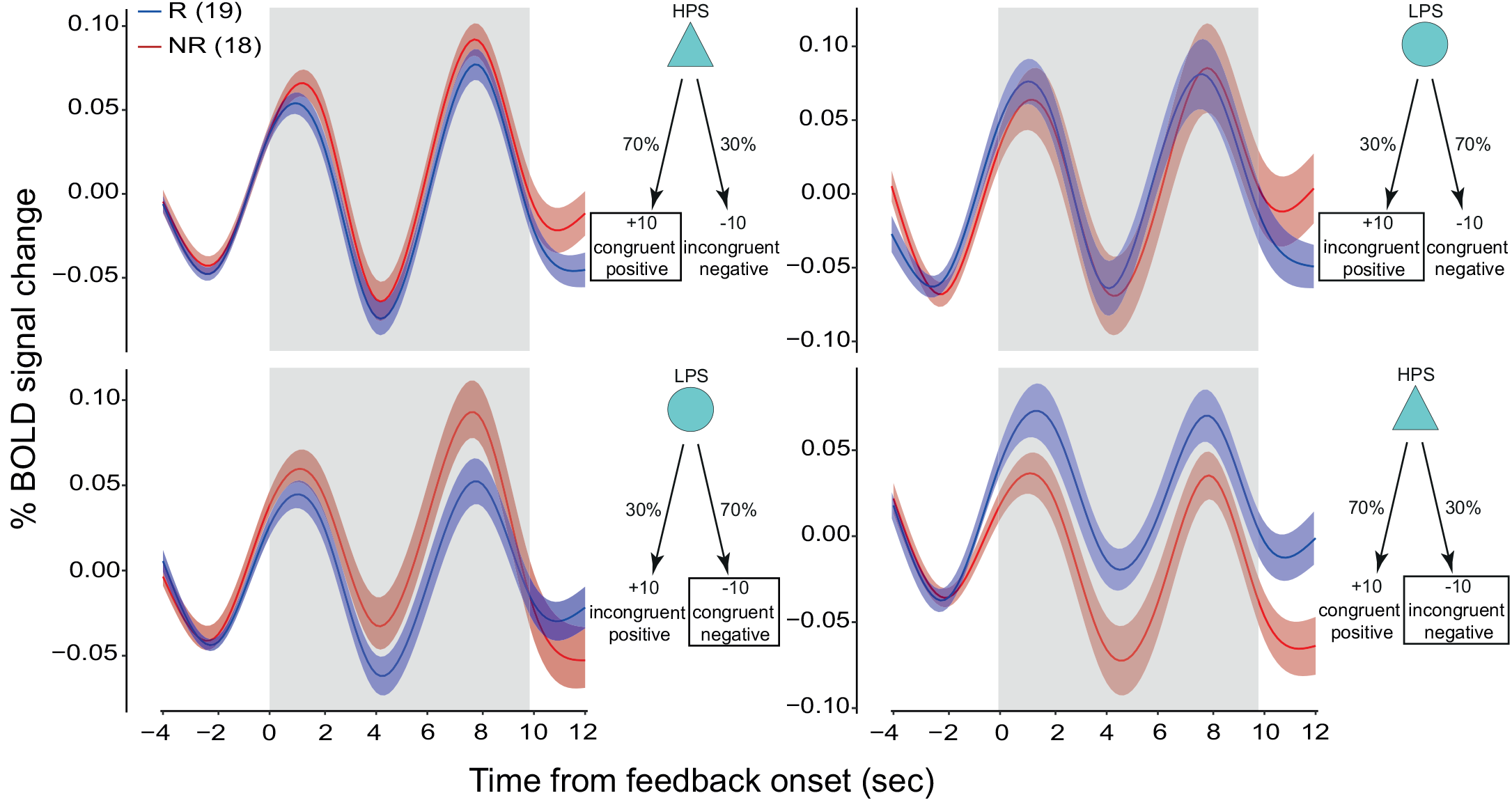
Pre-treatment time course of BOLD percent signal change in the dmPFC. Average feedback-locked BOLD percent signal change ± SEM (blue and red shading) in responders (blue; n=19) and non-responders (red; n=18) groups is plotted as a function of feedback valence and congruence. Averages are adjusted for pretreatment BDI-II score. We performed between-group comparisons over the temporal window shown by the grey shaded box. Responders exhibited comparatively lower BOLD activity during congruent negative trials (bottom left; F(1,33.01)=5.15, p=0.029) but comparatively higher BOLD activity during incongruent negative trials (bottom right; F(1,34.07)=8.87, p=0.005).

Without non-completers between-group differences remained significant only during incongruent negative trials (F_1,23.18_=4.99, p=0.03). No significant difference was detected regarding congruent negative feedback (F_1,23.02_=1.23, p=0.27) (Supplemental Figure S1). Moreover, we found a statistically significant negative correlation between treatment response magnitude and BOLD activity in incongruent negative trials (t_24_=-3.87, p<0.001) but not in congruent negative trials (t_24_=0.36, p=0.72) (Figure 7).

**Figure 7.**
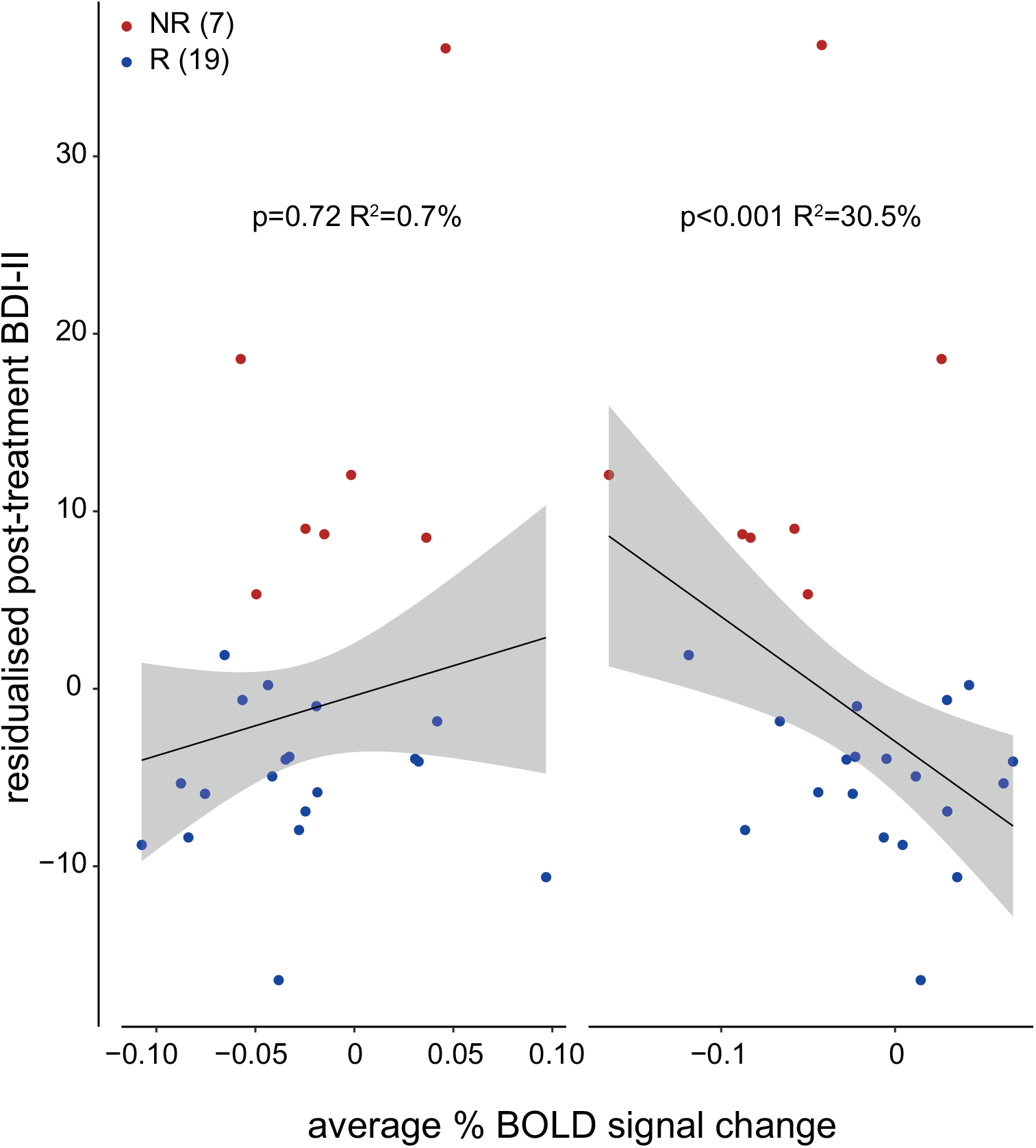
Relationship between BOLD percent signal change in the dmPFC and treatment response. Scatterplots show robust linear fit ± 95% CI (grey shading) of adjusted post-treatment BDI-II score against pre-treatment BOLD percent signal change in the dmPFC. Left: Congruent negative trials. Right: Incongruent negative trials. Post-treatment BDI-II score is adjusted for pre-treatment BDI-II score. Subject-wise pre-treatment BOLD percent signal change is averaged over a time window extending from 4 to 6 seconds after the onset of the outcome phase. Blue dots indicate responders (n=19) and red dots indicate non-responders (n=7).

## DISCUSSION

In the present study, to discriminate response to cCBT in depression we examined pre-treatment BOLD activity correlating with dynamic appraisal of probabilistic feedback during reinforcement learning. We found that greater BOLD activity in the dmPFC during incongruent negative trials and lower BOLD activity during congruent negative trials were associated with response to cCBT. Furthermore, we provided preliminary evidence that this pattern of BOLD activity may be moderating, rather than mediating, the effect of cCBT on clinical outcome (see Supplemental Results).

Crucially, although the focus of CBT in depression is centred in fostering adaptive reappraisal strategies (43), to the best of our knowledge no previous study has examined processing of probabilistic feedback as a function of response to CBT. Moreover, in spite of converging evidence that behavioural and neural responses to probabilistic feedback are abnormal in depression, the cognitive mechanisms underlying such impairment are still unknown and relatively unexplored. In this study we have addressed this knowledge gap by explicitly modelling the neurocomputational mechanisms of inference implicated in probabilistic reinforcement learning as a function of treatment response.

At a computational level we have capitalised on the notion that the model’s optimised fixed parameters capture a person’s typical mode of appraising incoming probabilistic information. We found that responders took greater account of previous feedback history than non-responders by means of greater smoothing over previous unsigned prediction errors. This operation helps averaging out noisy feedback from the trial-by-trial online computation of surprise and therefore makes inference more robust to random noisy fluctuations in the statistics of the environment. Somewhat complementary to this finding we found that responders were inclined to make smaller adjustments to the learning rate as a result of surprising outcomes. Again, this feature confers a relatively greater degree of robustness against noise to the inference process as it makes the temporal trajectory of the learning rate more stable.

On the whole, this specific information processing style fosters comparatively more farsighted and judicious decision-making in the responders group. CBT helps patients reframe extreme and dysfunctional negative thoughts in a more considered and balanced fashion (44). One fundamental requirement for CBT to be effective is the patient’s ability to actively engage in such work of cognitive restructuring. Depressed patients who are more adept at thoughtfully sieving through the barrage of noisy feedback information surrounding them exhibit a greater predisposition to critical thinking. This may translate in a greater ability to challenge maladaptive thinking patterns. Consistent with this line of reasoning is the prior finding that pre-treatment clinical ratings indicative of lower dysfunctional attitudes (as a result of less rigid and extreme thinking) predict better response to cCBT (12).

At the behavioural and neural level we have probed between-group differences in the online inference process underlying probabilistic learning, here quantified by a dynamic learning rate. Behaviourally model’s derived estimates of the dynamic learning rate did not discriminate response to cCBT. In contrast, BOLD activity did. Significant imaging but not behavioural results are not uncommon in cognitive neuroscience (45). One possible explanation is that underlying between-group differences are better captured by measures of neural activity. Additionally, lack of behavioural corroboration in our study may be due to the specific modelling approach employed here or simply to inadequate power. Crucially, our finding that neural, and not behavioural, measures of information processing discriminate response to cCBT underscores the translational potential of neuroimaging biomarkers in treatment prediction research in psychiatry.

We have shown that BOLD activity in the dmPFC underpins online appraisal of probabilistic feedback. Our cluster in the dmPFC largely overlaps with the anterior mid-cingulate cortex (aMCC). In a recent review of the functional roles ascribed to the aMCC Shackman et al. proposed that the core function of this region is to determine optimal choice behaviour under conditions of uncertainty (46). During probabilistic learning the differential evaluation of noisy and stochastic feedback is the main mechanism by which an agent tackles environmental uncertainty in the service of adaptive instrumental behaviour. In the computational framework employed here the dynamic learning rate encodes this flexible trial-wise weighting of information. Thus, our finding fits in with current theoretical accounts regarding the involvement of the aMCC in uncertainty resolution (46, 47).

In particular, we have shown that BOLD activity in the dmPFC denotes a tendency to assign comparatively smaller weight to negative, although still informative, feedback amongst responders. This may be symptomatic of a comparatively greater ability to maintain a positive outlook in spite of adverse events as the expected value of adverse events is more likely to be underestimated (48). Ultimately, a relatively greater capacity to lessen the detrimental impact that day-to-day setbacks can have on one’s own view of himself and the surrounding world may facilitate reframing of unhelpful thoughts and help boost response to CBT.

In contrast, during incongruent negative trials, BOLD activity in the dmPFC speaks to a relatively greater propensity to encode negative but less accurate feedback in the responders group and is thus expected to bias behavioural responses away from advantageous choices. Notably, this finding was robust to response bias and was further corroborated by a significant negative correlation between BOLD activity and post-treatment symptomatic improvement. Interestingly, our results concord with a previous report by Taylor Tavares et al. that lower BOLD activity in the dmPFC was associated with failure to disregard incongruent negative feedback (28).

At first glance, it would seem counterintuitive for such a pattern of pre-treatment BOLD activity to be associated with clinical improvement. Relatively greater underestimation of the value of generally advantageous choices may cause lack of engagement with positive states and ultimately engender an unduly negative view of the self and the world. One obvious interpretation is that this pattern of BOLD activity is the therapeutic target of CBT. However, this hypothesis is called into question by the observation that cross-session BOLD activity does not change within the responders group (see Supplemental Results). An alternative explanation is that a comparatively greater underestimation of advantageous choices serves the purpose of dampening down expectations on the amount of positive feedback that can be derived from those choices. This may render responders relatively more immune to the disappointment of high expectations not being met and thus buffer the deleterious consequences that negative, although somewhat misleading, events may have on mood.

In conclusion, in this study we have provided evidence supporting utility and feasibility of a neurocomputational approach to treatment response prediction in depression and, more in general, in mental health research. In particular, we have shown that computational and neural correlates of probabilistic reinforcement learning enable early discrimination of treatment response to cCBT in depression.

## ACKNOWLEDGEMENTS

The authors are grateful to Dr Rajeev Krishnadas for helpful discussion on the study, Dr John McLean for expert technical support and Prof Chris Williams for his help with the CBT-based intervention (http://llttf.com).

This work was supported by a Chief Scientist Office grant (PN09CP214) and the Dr Mortimer and Theresa Sackler Foundation. F.Q. is supported by an MRC- and MRF-funded clinical research training fellowship (PsySTAR). E.F. and M.G.P were supported by the Biotechnology and Biological Sciences Research Council (BBSRC; grant BB/J015393/2 to M.G.P.) and the Economic and Social Research Council (ESRC; grant ES/L012995/1 to M.G.P.).

---

**Supplemental Figure S1.**
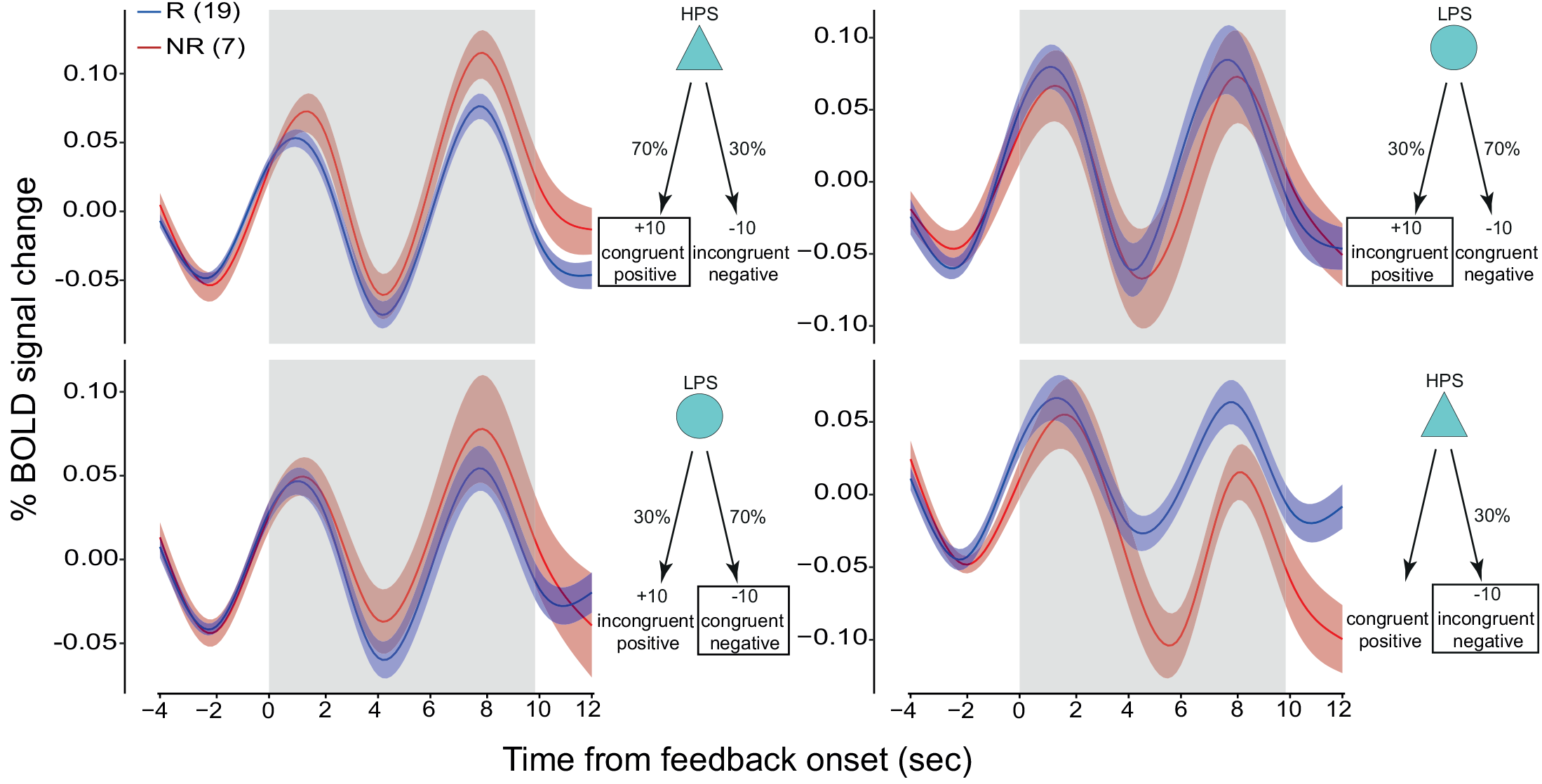
Pre-treatment time course of BOLD percent signal change in the dmPFC without non-completers. Average feedback-locked BOLD percent signal change ± SEM (blue and red shading) in responders (blue; n=19) and completers non-responders (red; n=7) groups is plotted as a function of feedback valence and congruence. Averages are adjusted for pre-treatment BDI-II score. We performed between-group comparisons over the temporal window shown by the grey shaded box. Between-group differences were robust to response bias during incongruent negative trials (bottom right; F(1,23.2)=4.99, p=0.03) but not during congruent negative trials (bottom left; F(1,22.6)=1.23, p=0.27).

